# Carbohydrate, glutathione, and polyamine metabolism are central to *Aspergillus flavus* oxidative stress responses over time

**DOI:** 10.1101/511170

**Authors:** Jake C. Fountain, Liming Yang, Manish K. Pandey, Prasad Bajaj, Danny Alexander, Sixue Chen, Robert C. Kemerait, Rajeev K. Varshney, Baozhu Guo

**Author notes:** Author e-mail addresses: J.C. Fountain; L. Yang; M. Pandey; P. Bajaj: p.; D. Alexander; S. Chen; R.C. Kemerait; R.K. Varshney; B. Guo. Corresponding Author: Dr. Baozhu Guo.

## Abstract

The primary and secondary metabolites of fungi are critical for adaptation to environmental stresses, host pathogenicity, competition with other microbes, and reproductive fitness. Drought-derived reactive oxygen species (ROS) have been shown to stimulate aflatoxin production and regulate development in *Aspergillus flavus*, and may function in signaling with host plants. Here, we have performed global, untargeted metabolomics to better understand the role of aflatoxin production in oxidative stress responses, and also explore isolate-specific oxidative stress responses over time. Two field isolates of *A. flavus*, AF13 and NRRL3357, possessing high and moderate aflatoxin production, respectively, were cultured in medium with and without supplementation with 15mM H_2_O_2_, and mycelia were collected following 4 and 7 days in culture for global metabolomics. Overall, 389 compounds were described in the analysis which were examined for differential accumulation. Significant differences were observed in both isolates in response to oxidative stress and when comparing sampling time points. The moderate aflatoxin-producing isolate, NRRL3357, showed extensive stimulation of antioxidant mechanisms and pathways including polyamines metabolism, glutathione metabolism, TCA cycle, and lipid metabolism while the highly aflatoxigenic isolate, AF13, showed a less vigorous response to stress. Carbohydrate pathway levels also imply that carbohydrate repression and starvation may influence metabolite accumulation at the later timepoint. Higher conidial oxidative stress tolerance and antioxidant capacity in AF13 compared to NRRL3357, inferred from their metabolomic profiles and growth curves over time, may be connected to aflatoxin production capability and aflatoxin-related antioxidant accumulation. The coincidence of several of the detected metabolites in H_2_O_2_-stressed *A. flavus* and drought-stressed hosts suggests their potential role in the interaction between these organisms and their use as markers/targets to enhance host resistance through biomarker selection or genetic engineering.

**Author Summary:** *Aspergillus flavus* is a fungal pathogen of several important crops including maize and peanut. This pathogen produces carcinogenic mycotoxins known as aflatoxins during infection of plant materials, and is particularly severe under drought stress conditions. This results in significant losses in crop value and poses a threat to food safety and security globally. To combat this, understanding how this fungus responds to environmental stresses related to drought can allow us to identify novel methods of mitigating aflatoxin contamination. Here, we analyzed the accumulation of a broad series of metabolites over time in two isolates of *A. flavus* with differing stress tolerance and aflatoxin production capabilities in response to drought-related oxidative stress. We identified several metabolites and mechanisms in *A. flavus* which allow it to cope with environmental oxidative stress and may influence aflatoxin production and fungal growth. These may serve as potential targets for selection in breeding programs for the development of new cultivars, or for alteration using genetic engineering approaches to mitigate excessive aflatoxin contamination under drought stress.

## Introduction

Abiotic stresses such as drought, heat, and osmotic stress have significant effects on the growth of plant pathogenic fungi, and can hinder their capability of infecting host plants. Drought stress in particular has been shown to have significant effects on both fungal pathogenicity and on host resistance to infection with some degree of specificity. For example, the growth of pathogenic fungi such as *Botrytis cinerea* causing gray mold, and *Oidium neolycopersici* causing powdery mildew on tomato are reduced or inhibited under drought stress (Achou et al. 2006; Ramegowda and Senthil-Kumar, 2015). Drought can also influence host metabolic composition and affect interactions with invading pathogens (Lecompte et al. 2017). The growth of microbes in soil environments along with their metabolic profiles and development can also be influenced by drought stress resulting in altered soil ecology and competition among microbes for limiting resources (Schimel et al. 2007). This shows the importance of metabolite accumulation in fungal environmental stress responses and pathogenicity, and the potential for abiotic stresses to regulate host plant immunity.

Members of the genus *Aspergillus* have been extensively studied using focused and untargeted metabolomics studies given their role as saprophytes in soil environments, their industrial applications, and their potential as human, animal, and plant pathogens. Examination of the metabolic responses of these fungi have been primarily focused on identifying metabolites involved in fungal growth and development, and the discovery of novel secondary metabolites encoded by silent, conserved gene clusters through both genomic prediction, and induced production using applied stressors or epigenetic modifying compounds (Albright et al. 2015 Bertrand et al. 2014; Brakhage, 2013; Scherlach and Hertweck, 2009). This has led to the identification of a number of metabolites with potential pharmaceutical applications, and several involved in the regulation of fungal biology (Amare and Keller, 2014; Calvo et al. 2002; Lopez et al. 2003). However, application of these techniques to study plant pathogenic species of *Aspergillus* have been limited.

An example of this is *Aspergillus flavus*, a facultative pathogen affecting crops such as maize and peanut which produces carcinogenic secondary metabolites termed aflatoxins. Annual losses can exceed $1 billion for US growers, particularly in regions susceptible to drought stress which has been shown to exacerbate aflatoxin contamination (Hill et al. 1983; Mitchell et al. 2016; Scully et al. 2009). Recent examination of developing maize kernels under drought stress also showed that accumulation of polyunsaturated fatty acids, and simple sugars along with decreases in antioxidants such as polyamines occurred in inbred lines sensitive to drought stress and susceptible to aflatoxin contamination (Yang et al. 2018). The same study also showed a greater accumulation of reactive oxygen species (ROS), specifically hydrogen peroxide (H_2_O_2_), in kernels of the drought sensitive line compared to the drought tolerant line under drought suggesting a correlation between drought tolerance and both ROS accumulation and aflatoxin contamination. Therefore, investigating this correlation between both matrix composition and ROS accumulation with aflatoxin production in *A. flavus* may provide insights into mitigating drought-induced contamination.

Matrix composition has been shown to heavily influence both *A. flavus* growth and aflatoxin production. For example, carbon sources have been found to have a significant effect on aflatoxin production *in vitro* with simple sugars being able to support aflatoxin production by *A. flavus* and *A. parasiticus* while other carbon sources such as peptone can inhibit aflatoxin production in a concentration-dependent manner (Fountain et al. 2015; Yan et al. 2012). Carbon source and availability has also been shown to influence conidiation in *A. flavus* (Fountain et al. 2015). In addition, the accumulation of lipid compounds such as unsaturated fatty acids, and oxylipins in host tissues have been demonstrated to influence aflatoxin production (Burow et al. 1997; Gao et al. 2009; Xue et al. 2003; Zerinque et al. 1996).

During *in vitro* experiments, the same ROS detected by Yang et al. (2018) to accumulate in maize kernels under drought have also been shown to stimulate the production of aflatoxin in both *A. flavus* and *A. parasiticus*, and aflatoxin precursors in *A. nidulans* (Grintzalis et al. 2014; Jayashree and Subramanyam, 2000; Narasaiah et al. 2006; Yin et al. 2013). Variation in oxidative stress tolerance has also been observed among field and mutant isolates of *A. flavus* with isolates exhibiting greater aflatoxin production and more later-stage precursor production tending to tolerate greater levels of oxidative stress compared to less toxigenic or atoxigenic ones (Fountain et al. 2015; Roze et al. 2015). Such variation in stress tolerance and growth patterns may also be characteristic of differences in vegetative compatibility groups (VCGs) which have been shown to vary in host pathogenicity, and competitive ability with other isolates for environmental nutrients (Mehl and Cotty, 2010, 2013). Also, ROS function in reproductive signaling in *Aspergillus spp.* with oxidative responses being closely interconnected with the regulation of reproductive development (Roze et al. 2011). Further, oxidative stress results in extensive metabolic profile alterations to fungi with regards to primary metabolism and antioxidant mechanisms following induction by either ROS or ROS-generating compound application (Sobon et al. 2018; Xu et al. 2018; Zheng et al. 2015).

Previous experimentation examining the oxidative stress responses of field isolates of *A. flavus* with different levels of aflatoxin production and stress tolerance by our group have shown a high degree of variability among isolates in overall strategies to remediate stress at both the transcript and protein levels (Fountain et al. 2016a, 2016b, 2018). These studies suggested that highly toxigenic isolates may exhibit earlier, more effective oxidative stress remediation mechanisms compared to less toxigenic or atoxigenic isolates. Transcripts and proteins involved in antioxidant protection, carbohydrate metabolism, microbial competitiveness, reproductive development, and the production of other secondary metabolites such as kojic acid and aflatrem were among those differentially expressed in response to oxidative stress. Differences in isolate-specific oxidative stress responses were also proposed to be due to resource allocation and the regulation of primary and secondary metabolic pathways to mitigate oxidative damage. While these studies provided an extensive overview of transcript and protein-level responses to oxidative stress, they are not fully capable of characterizing changes in final biochemical product levels, and resource allocation over time. Therefore, the objectives of this study were: 1) to identify differentially accumulating metabolites over time to explain isolate-to-isolate variability in oxidative stress responses; 2) to identify metabolic responses that begin to explain the relationship between oxidative stress and exacerbated aflatoxin production; and 3) to identify the metabolites that correspond to host drought responses with potential use in improving host resistance through selection or biotechnology. To accomplish this, we performed a global, untargeted metabolomics analysis of two field isolates of *A. flavus* with different levels of aflatoxin production and their response to oxidative stress over time.

## Results

### Effects of oxidative stress on isolate growth rates

Two isolates of *A. flavus*, AF13 and NRRL3357, which were previously observed to possess relatively high (up to 35mM H_2_O_2_) and moderate (up to 20mM H_2_O_2_) levels of oxidative stress tolerance and aflatoxin production, respectively (Fountain et al. 2015), were selected for this study. The isolate AF13 is a high aflatoxin producing L-strain (sclerotia size >400µm) with a MAT1-2 mating type belonging to the YV-13 vegetative compatibility group (VCG) and relatively high tolerance to oxidative stress (Cotty, 1989; Ehrlich et al. 2007; Fountain et al. 2015). The isolate NRRL3357 is a moderately high aflatoxin producing L-strain with a MAT 1-1 mating type with no currently defined VCG, and moderate tolerance to oxidative stress (Chang et al. 2012; Fountain et al. 2015). These isolates were examined for conidial oxidative stress tolerance and the effect of oxidative stress on growth rates. Increasing levels of stress caused significant delays in the initial detection (T_i_) of isolate growth for both isolates, but to a greater extent in NRRL3357 compared to AF13 at both inoculum levels (Figure 1, Table S1). For AF13, significant growth delays were observed beginning at 10mM H_2_O_2_ and increasing up to 25mM where growth was completely suppressed at 20,000 conidia/mL (Figure 1A) but not at 80,000 conidia/mL (Figure 1C). However, growth was completely inhibited at 30mM H_2_O_2_ even at the higher inoculum concentration (Table S1). For NRRL3357, significant delays in growth were also observed at 10mM H_2_O_2_ while 15mM was completely inhibitory of growth at 20,000 conidia/mL (Figure 1B) but not at 80,000 conidia/mL (Figure 1D). Growth was also completely inhibited at 20mM H_2_O_2_ at the higher inoculum concentration (Table S1). These inhibitory concentrations of H_2_O_2_ observed for each isolate were 5 – 10mM less than observed when the isolates were cultured in H_2_O_2_ amended YES medium in Erlenmeyer flasks with cotton plugs (Fountain et al. 2015).

**Figure 1.**
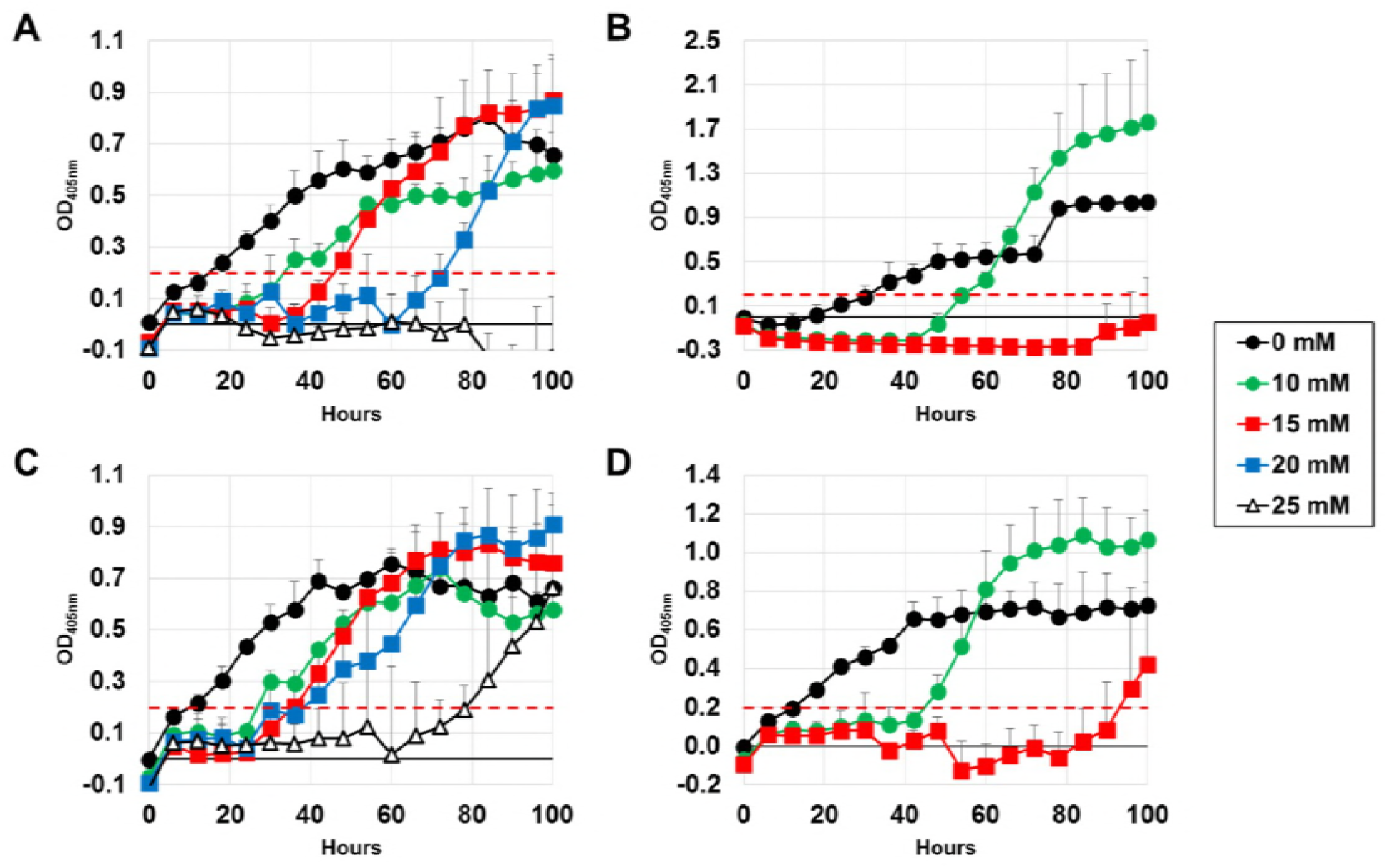
Growth curve analysis of *Aspergillus flavus* isolates AF13 and NRRL3357 under increasing oxidative stress and conidial concentration. The growth of AF13 (A, C) and NRRL3357 (B, D) were examined under increasing H_2_O_2_ concentrations in YES medium inoculated with either 2.0 x 10^4^ (A, B) or 8.0 x 10^4^ conidia/mL (C, D) by monitoring absorbance at 405 nm over 100 hours. A threshold of 0.2 was selected for growth initiation timing which corresponded with linear phase initiation for both isolates in most conditions and is indicated by the dashed red line. Error bars represent standard deviation. No growth was detected at H_2_O_2_ concentrations >15mM in NRRL3357 with earlier growth initiation detected at higher conidia concentrations.

### Differential metabolic alterations in response to oxidative stress over time

Two field isolates of *A. flavus* were selected for global, untargeted metabolomics analysis using an UPLC-MS/MS approach to examine their responses to drought-related, H_2_O_2_-derived oxidative stress over time. This metabolomics analysis identified 389 distinct metabolites. Functional classification for the detected metabolites was performed based on the Kyoto Encyclopedia of Genes and Genomes (KEGG) database (Kanehisa and Goto, 2000). These metabolites were grouped into nine super pathways with a majority of metabolites being classified as either amino acids (163), lipids (84), nucleotides (54), or carbohydrates (43). These super pathways were further divided into 47 sub-pathways which are described in Table S2.

Welch’s two-sample t-test was used for differential accumulation analyses to identify metabolites significantly different between oxidative stress treatments, between isolates, or over time (Table 1). Data normalization using DNA or protein content was found to introduce possible skewing in time and isolate effects on metabolite accumulation. Therefore sample mass per unit volume of extraction solvent was used for normalization and these data were used for analysis and interpretation. Both protein and non-protein normalized datasets, and raw data are included in Supplemental File 1. When comparing between stress treatments, AF13 showed 111 and 47 metabolites which differentially accumulated at 4 and 7 DAI, respectively. Of these, 27 and 64 metabolites were significantly increased and decreased, respectively, in abundance at 4 DAI, and 34 and 13 were increased and decreased in abundance, respectively, at 7 DAI. For NRRL3357, 223 and 90 metabolites were differentially accumulated at 4 and 7 DAI, respectively, in response to stress. Of these, 90 and 133 were significantly increased and decreased, respectively, at 4 DAI, and 65 and 25 were increased and decreased in abundance, respectively, at 7 DAI. Time was a highly significant influence on metabolite accumulation with AF13 showing 257 metabolites with significant differences in abundance between 4 and 7 DAI without H_2_O_2_treatment and 268 with H_2_O_2_ treatment. In NRRL3357, this was also apparent with 243 and 261 metabolites being significantly altered in abundance between 4 and 7 DAI either with or without H_2_O_2_ treatment, respectively. Comparisons between the isolates are more likely to reflect genetic differences rather than stress response, but more stark differences in numbers of differentially accumulating metabolites could be observed between AF13 and NRRL3357 at 4 DAI regardless of H_2_O_2_ treatment.

**Table 1.**
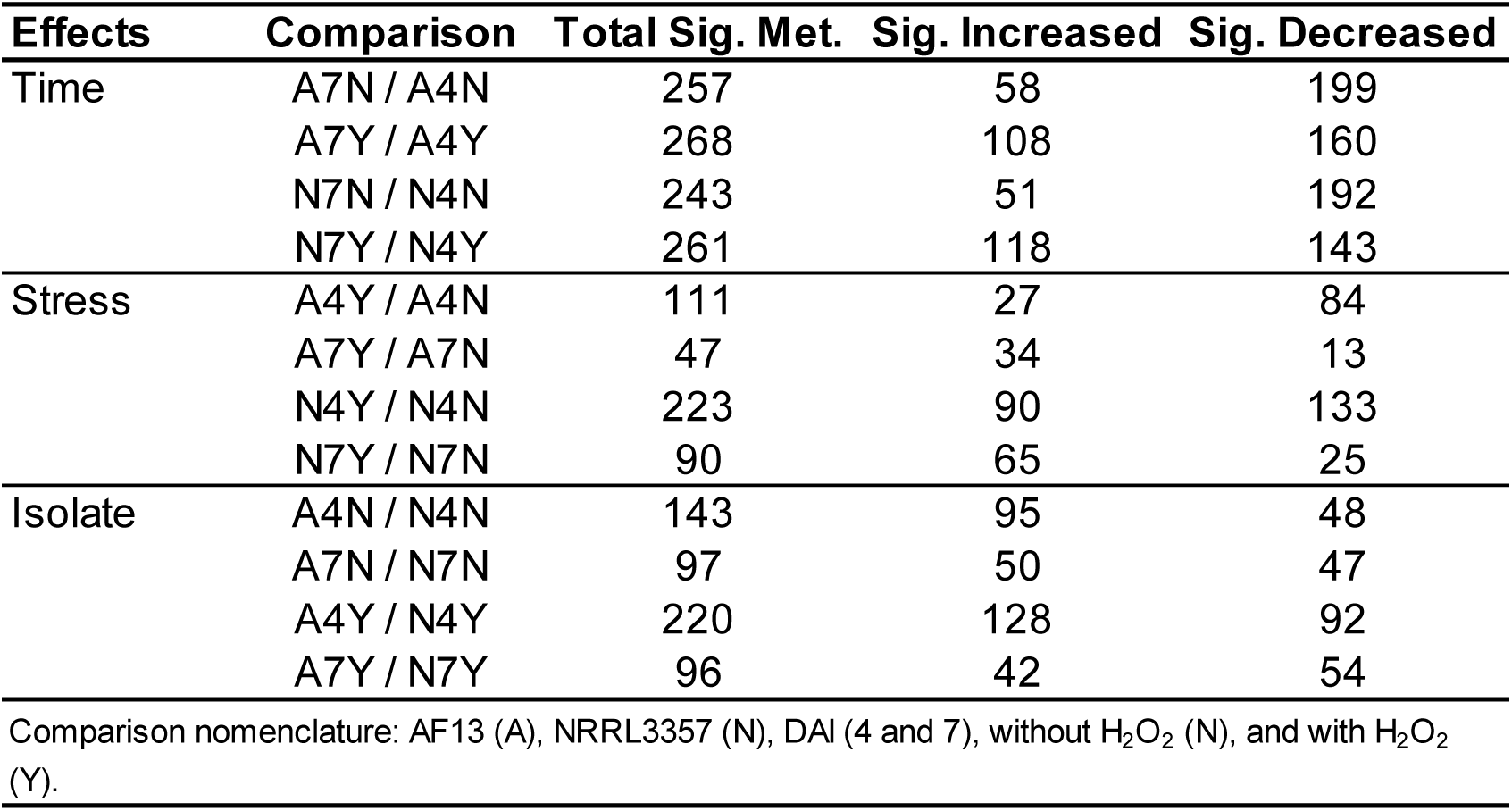
Numbers of significantly, differentially accumulating metabolites.

| Effects | Comparison | Total Sig. Met. | Sig. Increased | Sig. Decreased |
| --- | --- | --- | --- | --- |
| Time | A7N / A4N | 257 | 58 | 199 |
|  | A7Y / A4Y | 268 | 108 | 160 |
|  | N7N / N4N | 243 | 51 | 192 |
|  | N7Y / N4Y | 261 | 118 | 143 |
| Stress | A4Y / A4N | 111 | 27 | 84 |
|  | A7Y / A7N | 47 | 34 | 13 |
|  | N4Y / N4N | 223 | 90 | 133 |
|  | N7Y / N7N | 90 | 65 | 25 |
| Isolate | A4N / N4N | 143 | 95 | 48 |
|  | A7N / N7N | 97 | 50 | 47 |
|  | A4Y / N4Y | 220 | 128 | 92 |
|  | A7Y / N7Y | 96 | 42 | 54 |

Comparison nomenclature: AF13 (A), NRRL3357 (N), DAI (4 and 7), without H<sub>2</sub>O<sub>2</sub> (N), and with H<sub>2</sub>O<sub>2</sub> (Y).

These differences in metabolite accumulation were also observed in principal components analyses (Figure 2). The first component was dominated primarily by time effects reflecting significant differences between the 4 and 7 DAI time points. Significant stress effects could also be observed between the isolates with more stark differences observed at 4 DAI. Samples from 7 DAI did not segregate into distinct clusters as seen in samples from 4 DAI. A higher degree of variability between biological replicates was also observed in the 7 DAI samples compared to 4 DAI.

**Figure 2.**
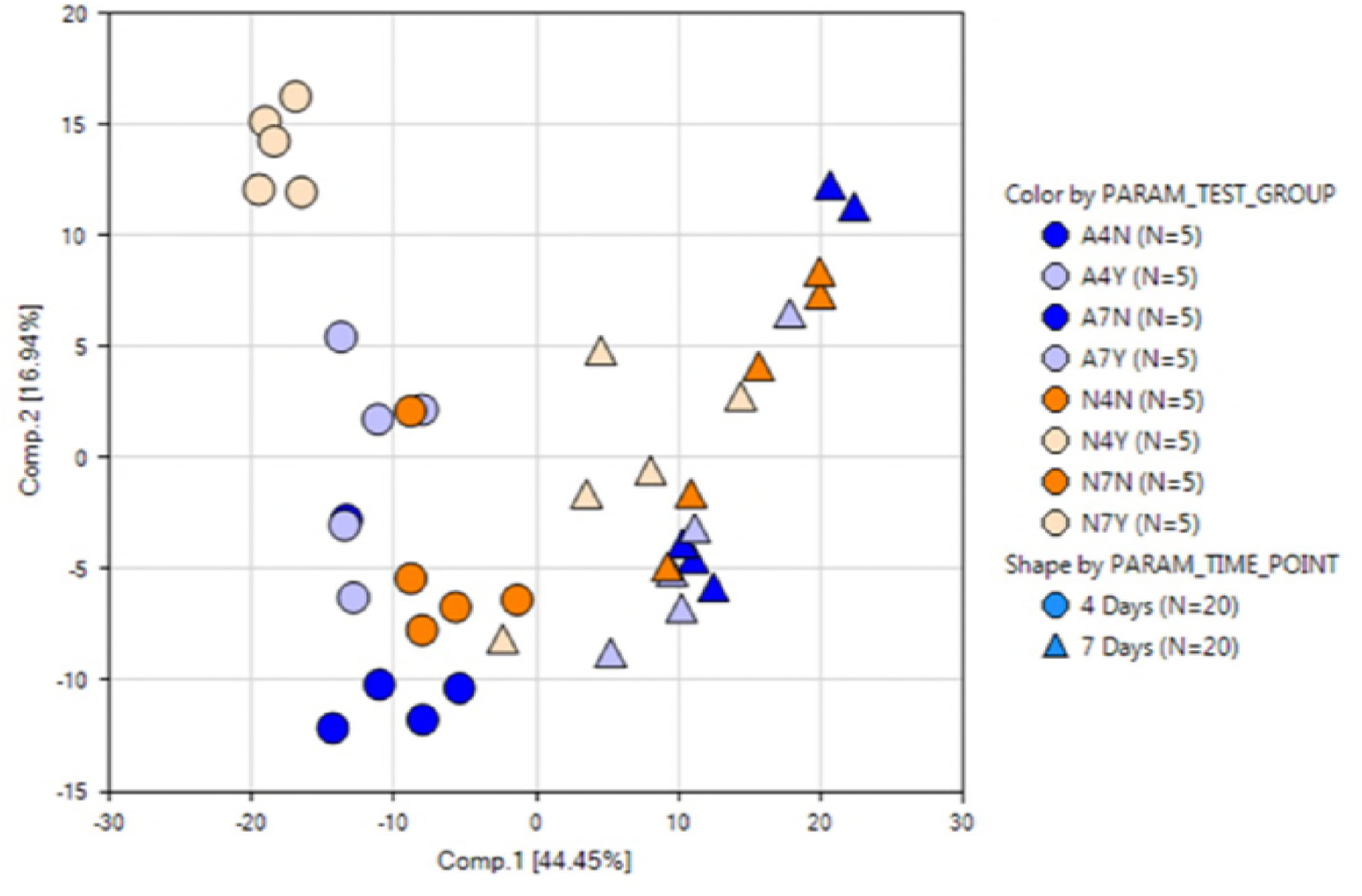
Principal components analysis (PCA) of metabolite accumulation. A4N, A4Y, A7N, and A7Y refer to AF13 at 4 and 7 DAI with and without 15mM H_2_O_2_ treatment. N4N, N4Y, N7N, and N7Y refer to the same for NRRL3357. Dark blue points correspond with AF13 with no stress and light blue points refer to AF13 with stress. Orange points correspond with NRRL3357 with no stress and light orange points refer to NRRL3357 with stress. Circles represent samples at 4 DAI and triangles represent samples at 7 DAI.

### Carbohydrate metabolic responses to oxidative stress

Significant variation in carbohydrate metabolite accumulation was observed in response to oxidative stress in both AF13 and NRRL3357. AF13 showed significant changes in glycolytic compounds glucose and pyruvate with significant decreases (p < 0.05) in both compounds at 4 DAI in response to stress with glucose and fructose levels showing marginally significant increases at 7 DAI (p < 0.10; Figure 3). NRRL3357 showed significant decreases in both glucose and pyruvate at 4 DAI in response to stress with a significant decrease in pyruvate also detected at 7 DAI. Fructose levels in NRRL3357 were also increased at both time points in response to stress (Figure 3). Time effects showed that pyruvate accumulated in NRRL3357 over time regardless of H_2_O_2_ treatment, and glucose and fructose were significantly decreased over time with reductions in glucose only seen in non-stressed samples (Supplemental File 1).

**Figure 3.**
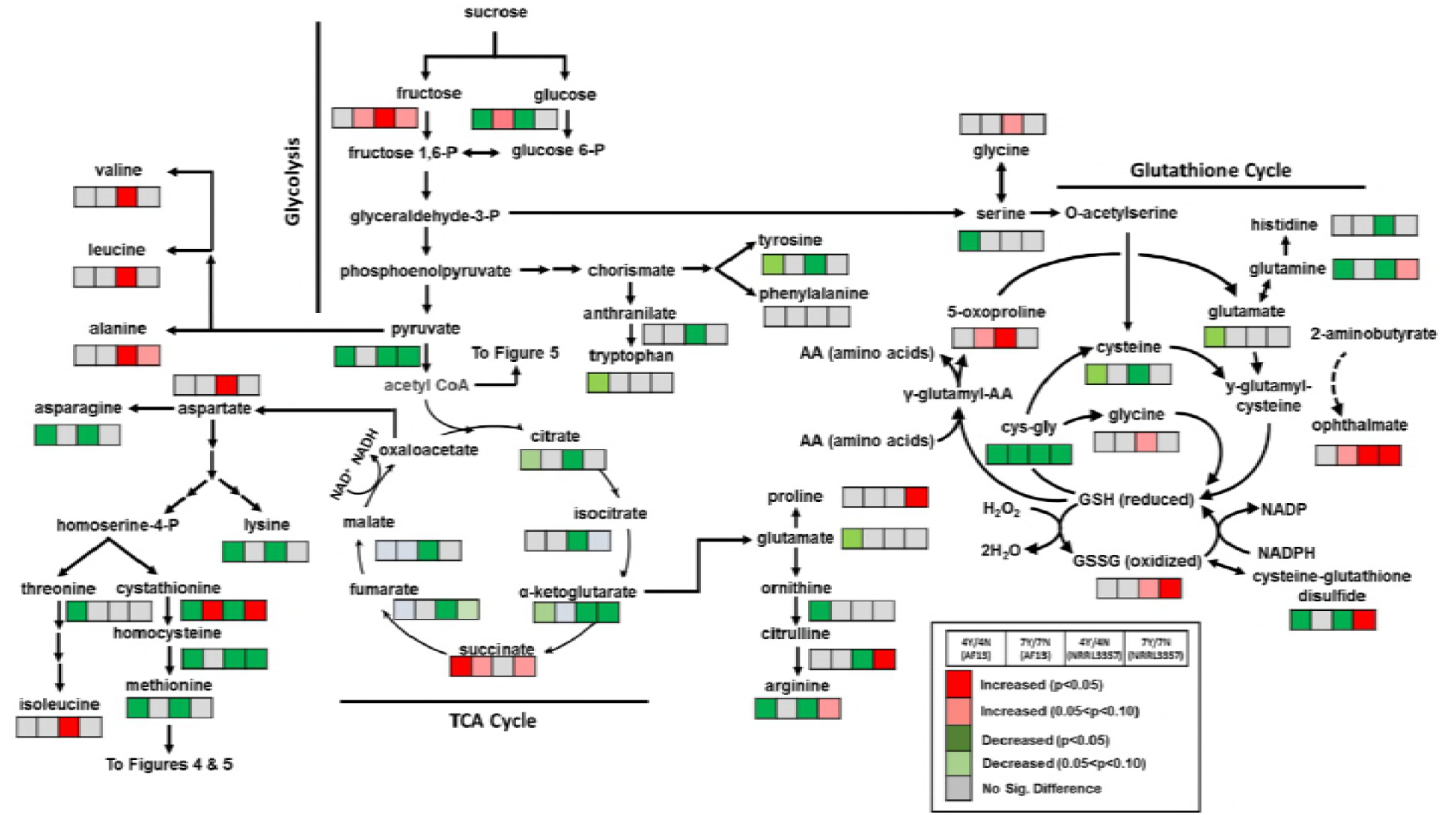
Differential accumulation of compounds involved in carbohydrate metabolism, glutathione metabolism, and amino acid biosynthesis. Heatmaps located at each metabolite represent the changes in metabolite accumulation in response to oxidative stress in AF13 and NRRL3357 at 4 and 7 DAI. Red and green indicate significant increases and decreases in metabolite levels, respectively (p < 0.05). Light red and light green indicate marginally significant increases and decreases in metabolite levels, respectively (0.05 < p < 0.10). Grey represents no significant changes.

Intermediates in the tricarboxylic acid (TCA) cycle were also significantly affected by oxidative stress. NRRL3357 showed significant reductions, particularly at 4 DAI, in citrate, isocitrate, alpha-ketoglutarate, fumarate, and malate in response to oxidative stress (Figure 3). Conversely, AF13 showed no significant changes in TCA intermediate levels in response to oxidative stress with the exception of a significant increase in succinate at 4 DAI. These compounds were, however, seen to generally accumulate over time in the stressed samples when comparing time points (Supplemental File 1).

In addition to these pathways, AF13 and NRRL3357 showed significant reductions in trehalose, arabitol, and xylitol in response to oxidative stress at 4 DAI with less significant decreases or no significant differences being observed in response to stress at 7 DAI (Supplemental File 1). Additional metabolic products of arabinose and xylinose, arabinate and xylinate were increased in accumulation in response to stress in both isolates and time points (Supplemental File 1). Increases in amino sugars were also observed in both isolates, particularly at 4 DAI in response to stress (Supplemental File 1).

### Amino acid metabolic responses to oxidative stress

Significant changes in the accumulation of amino acids and their derivatives were observed in both isolates in response to oxidative stress over time. Changes in primary amino acids were proportional to changes in their precursors with more significant changes occurring in NRRL3357 compared to AF13 (Figure 3; Supplemental File 1). In particular, changes in aromatic amino acid precursors in the tryptophan and histidine pathways were observed in NRRL3357 in response to oxidative stress although the levels of tryptophan and histidine were unchanged or reduced, respectively, in the same conditions (Supplemental File 1). In addition, the tryptophan derivative kynurenine was increased in NRRL3357 at both time points in response to oxidative stress, but not in AF13 (Supplemental File 1). Proline levels were also increased in NRRL3357 at 7 DAI in response to stress (Supplemental File 1). Among the amino acid derivatives, those involved in glutathione, polyamine, and sulfur metabolism were among the most differentially accumulating in response to oxidative stress.

Glutathione metabolism was significantly regulated in both isolates but to a greater extent in NRRL3357 compared to AF13 (Figure 3). Significant increases in 5-oxoproline, ophtalmate, oxidized glutathione (GSSH), and cysteine-glutathione disulfide were observed in NRRL3357 in response to increasing stress (Figure 3). AF13 showed marginally significant (p < 0.10) increases in accumulation of only 5-oxoproline and ophtalmate were see at 7 DAI in response to stress. Direct comparison of levels between these isolates showed that AF13 accumulated significantly greater levels of GSSH and cysteine-glutathione disulfide at 4 DAI compared to NRRL3357 in the absence of oxidative stress, and equivalent and greater levels, respectively, of each when under oxidative stress (Supplemental File 1). When comparing time point measurements, 5-oxoproline, ophtalmate, and GSSG showed significant reductions in accumulation in both isolates and treatments (Supplemental File 1). Significant changes were also found among the gamma-glutamyl amino acids which were significantly reduced in AF13 at 4 DAI in response to stress, but tended to be either unchanged or increased in accumulation in NRRL3357 in response to stress (Figure 3).

In addition to glutathione, other sulfur-containing amino acids and their metabolites were significantly regulated in response to oxidative stress (Figure 4). Significant reductions in methionine levels were observed in both isolates at 4 DAI in response to oxidative stress. S-adenosylmethionine (SAM), an important signaling compound, was also significantly regulated in response to oxidative stress showing increasing accumulation at 7 DAI in both isolates, and a significant decrease at 4 DAI in NRRL3357 (Figure 4). 5-methylthioadenesine (MTA) also exhibited a similar pattern of accumulation to SAM. In addition to methionine derivatives, cysteine also serves as a precursor to the antioxidant compound taurine which was significantly increased in both isolates at 4 DAI and in AF13 at 7 DAI in response to oxidative stress. A taurine precursor, 3-sulfo-L-alanine, was also significantly increased in both isolates and time points in response to oxidative stress (Figure 4).

**Figure 4.**
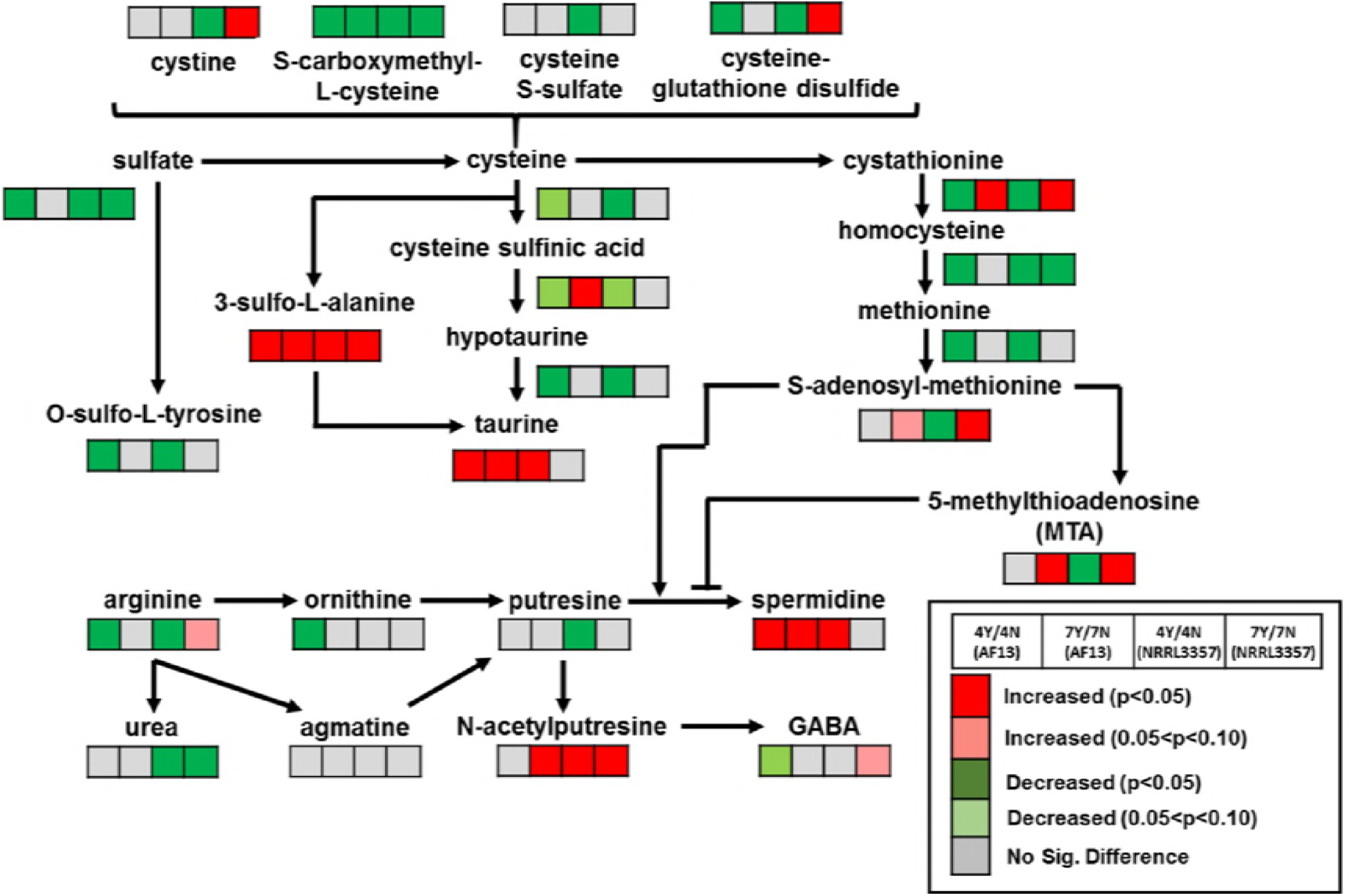
Differential accumulation of compounds involved in polyamine and sulfur metabolism. Heatmaps located at each metabolite represent the changes in metabolite accumulation in response to oxidative stress in AF13 and NRRL3357 at 4 and 7 DAI. Red and green indicate significant increases and decreases in metabolite levels, respectively (p < 0.05). Light red and light green indicate marginally significant increases and decreases in metabolite levels, respectively (0.05 < p < 0.10). Grey represents no significant changes.

Polyamine metabolites were also significantly regulated in response to oxidative stress in both isolates. Ornithine showed significant reduction in AF13 at 4DAI while putresine showed the same in NRRL3357 in response to oxidative stress while the immediate precursor to ornithine, N-alpha-acetylornithine, was increased in both isolates at 4 DAI (Figure 4; Supplemental File 1). These compounds are precursors to both spermidine and N-acetylputresine which showed significant increases in both isolates in response to oxidative stress. N-acetylputresine is a part of butanoate metabolism and used for the biosynthesis of gamma-aminobutanoate (GABA) which showed marginally significant changes in abundance in response to oxidative stress (Figure 4).

### Fatty acid metabolic responses to oxidative stress

Several fatty acids and their derivatives were also significantly regulated in response to H_2_O_2_-stress over time. Significant regulation of saturated and mono- and poly-unsaturated fatty acid accumulation were primarily observed in NRRL3357 in response to stress (Figure 5). Significant increases in the saturated fatty acids pentadecanoic acid (15:0) and heptadecanoic acid (17:0) were seen at 7 and 4 DAI, respectively, in NRRL3357 in response to stress (Figure 5). Similarly, significant increases in several unsaturated fatty acids were also seen in NRRL3357 (Figure 5). While no significant regulation of these fatty acids was seen in AF13 in response to oxidative stress within each time point, significant depletion of these fatty acids was observed in both isolates over time with or without the presence of oxidative stress (Supplemental File 1).

**Figure 5.**
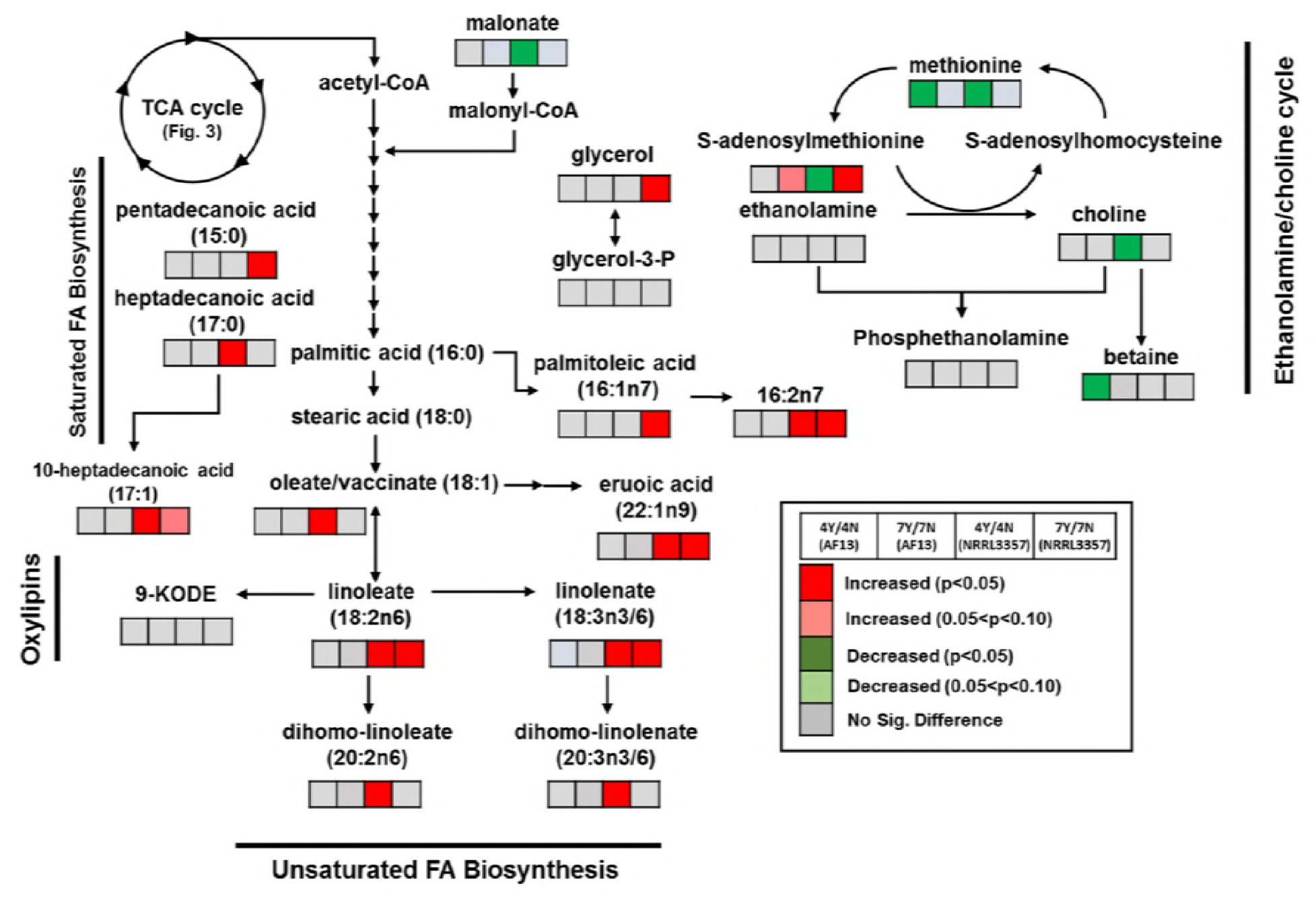
Differential accumulation of compounds involved in lipid metabolism. Heatmaps located at each metabolite represent the changes in metabolite accumulation in response to oxidative stress in AF13 and NRRL3357 at 4 and 7 DAI. Red and green indicate significant increases and decreases in metabolite levels, respectively (p < 0.05). Light red and light green indicate marginally significant increases and decreases in metabolite levels, respectively (0.05 < p < 0.10). Grey represents no significant changes.

Other fatty acid derivatives were also found to differentially accumulate in the isolates under oxidative stress. In AF13, betaine, an ethanolamine derivative, was significantly decreased under oxidative stress at 4 DAI (Figure 5). Several phospholipids such as glycerophosphoglycerol were also found to be differentially accumulating in response to oxidative stress in both isolates (Supplemental File 1). Ergosterol levels were found to be significantly decreased under stress in AF13 at 7 DAI and in NRRL3357 at 4 DAI. There was no significant change in ergosterol levels over time in either treatment, but AF13 accumulated significantly more than NRRL3357 at 4 DAI in both treatments, while at 7 DAI AF13 was found to have more only in the non-stressed control (Supplemental File 1).

### Other compounds regulated in response to oxidative stress

In addition to amino acids, carbohydrates, and lipids, other classes of compounds were found to differentially accumulate in response to increasing oxidative stress over time in both isolates. Among cofactors and electron carriers, carnitine and related metabolites were significantly reduced in both isolates at 4 DAI in response to stress, but showed significant increases over time under stress and was present in higher concentrations in AF13 compared to NRRL3357 (Supplemental File 1). Several B vitamins with potential antioxidant activity were also regulated in response to stress including thiamin (B1), riboflavin (B2), and pyridoxine (B6) (Supplemental File 1). Several nucleotide derivatives were also differentially accumulated in response to stress such as adenosine 5’-monophosphate (AMP) which was significantly increased in AF13 at 4 DAI in response to stress, but was significantly depleted in both isolates and treatments over time (Supplemental File 1). Terpenoid and isoterpenoid precursors were also found to differentially accumulate under stress with mevalonate along with its immediate precursor, 3-hydroxy-3-methylglutarate, and its lactone form, mevalonolactone, showing significant increases in response to stress in AF13 at 7 DAI and in NRRL3357 at 4 and 7 DAI (Supplemental File 1).

## Discussion

Drought stress is one of the primary factors contributing to the exacerbation of pre-harvest aflatoxin contamination in the field. Drought stress has been shown to significantly alter the metabolic composition of maize kernels during earlier stages of development resulting in increased levels of free simple sugars, oxylipins, free fatty acids, and signaling compounds including ROS including H_2_O_2_, superoxide (O_2_ ^-^), and hydroxyl ions (OH^.-^) (Yang et al. 2015, 2018). These ROS have also been found to stimulate and be required for aflatoxin production (Jayashree and Subramanyam, 2000). These observations served as the impetus to investigate responses of *A. flavus* isolates with varying levels of aflatoxin production to drought-related oxidative stress and the metabolite level over time.

Aflatoxin production capability has been previously correlated with *A. flavus* isolate oxidative stress tolerance (Fountain et al. 2015; Roze et al. 2015). When examining the growth rates and behavior of *A. flavus* isolates under oxidative stress AF13, a highly toxigenic isolate, was found to exhibit higher levels of oxidative stress tolerance and growth under stress compared to NRRL3357, a moderately high toxigenic isolate (Figure 1). Aflatoxin production and the reactions in the biosynthetic pathway are suspected to result in increased conidial oxidative stress tolerance due to stimulating additional antioxidant enzyme production, or through the consumption of ROS during production (Fountain et al. 2016a; Roze et al. 2015). Given this, conidial antioxidant enzyme activity may have contributed here. To further examine this, growth curve analyses was performed with different inoculum concentrations, and showed that growth for both isolates occurred at elevated H_2_O_2_ levels when the inoculum was increased from 20,000 conidia/mL as described by Meletiadis et al. (2001) to 80,000 conidia/mL used here as inoculum for cultures used for metabolomics analysis. Interestingly, even with the increased conidia concentration, observed stress tolerance remained approximately 5 mM less than the maximum observed in the previous study. While this may be an artifact of performing the assay in a sealed microplate with potentially limited oxygen availability, the overall trend was consistent with previous observations of each isolate’s tolerance to oxidative stress (Clevstrom et al. 1983; Fountain et al. 2015; Jayashree and Subramanyam, 2000).

When examining overall metabolite accumulation patterns, NRRL3357 displayed approximately double the number of differentially accumulating metabolites compared to AF13 in response to oxidative stress with both isolates exhibiting greater numbers at 4DAI compared to 7 DAI (Table 1). This pattern mirrors observed numbers and functional classifications of differentially expressed transcripts and proteins for these isolates in response to similar levels of oxidative stress in our previous transcriptome and proteome studies (Fountain et al. 2016a, 2016b, 2018). Here, significant differences in metabolite accumulation were detected within and between time points in both isolates, and sampling time was one of the major grouping factors in the PCA analysis (Table 1; Figures 2 and S1). These time course influences may be due to differences in isolate growth patterns and, presumably, timing and vigor of oxidative stress remediation mechanisms. As indicated by the growth curve analysis (Figure 1; Table S1), earlier initiation of growth in AF13 compared to NRRL3357 may be the result of earlier, more vigorous lag phase or conidial oxidative stress remediation processes. Therefore, sampling at 4 DAI for both isolates would describe actively growing and responding tissues while sampling at 7 DAI would describe stationary state responses in AF13 having already remediated the majority of oxidative stress while NRRL3357 would still be actively growing and responding to stress.

Examining the isolate-specific responses to oxidative stress, there were significant differences in carbohydrate accumulation. Both glycolysis and TCA cycle intermediates were significantly altered in accumulation in response to oxidative stress in these isolates, but to differing degrees. NRRL3357 displayed increased demand for TCA intermediates showing significant decreases in most quantified metabolites in the cycle while AF13 showed no significant differences (Figure 3). These compounds have been shown to provide some antioxidant benefit when supplemented to cultured neuronal cells (Sawa et al. 2017), though a more likely explanation is the use of these compounds in the synthesis of amino acids and/or their derivatives involved in oxidative stress remediation. Increases in glucose and fructose under stress in both isolates may also be reflective of higher levels of metabolic demand for simple sugars, and the beginnings of carbon starvation leading to gluconeogenesis (Dijkema et al. 1985; Lima et al. 2014), particularly at 7 DAI (Figure 3).

When examining time course effects, the accumulation of these compounds in stressed samples may also be indicative on increased energy demand and the need to maintain redox homeostasis through the generation of reduced coenzymes for oxidative phosphorylation such as NADH and NADPH (Kapoor et al. 2015). Significant reductions in their accumulation in NRRL3357 under stress (Figure 3) could, therefore, partially explain the reduced growth rate, and observed ongoing stress responses compared to AF13. Also of interest, the pentose phosphate pathway has been shown to be involved in oxidative stress responses in yeast, and amino sugars such as ribonate are also involved in the generation of reduced coenzymes used for redox homeostasis (Campbell et al. 2016; Juhnke et al. 1996). These reduced coenzymes, particularly NADPH, are also critical for the activity of polyketide synthases which may also impact aflatoxin production levels under oxidative stress (Huang et al. 2009; Kletzien et al. 1994; Shih and Marth, 1974).

Changes in amino acid metabolite levels appeared to form the basis of a majority of the oxidative stress remediation processes employed by these isolates constituting the bulk of directly antioxidant compounds and mechanisms. Amino acids such as proline have been previously shown to be involved in osmotic, drought, and oxidative stress tolerance in fungi and plants and were increased in abundance in NRRL3357 (Chen and Dickman, 2005; Szabados and Savoure, 2009). Also in NRRL3357, the tryptophan derivative kynurenine was increased and may also contribute to stress remediation. Disruption of kynurenine 3-monooxygenase, a central enzyme in kynurenine metabolism, in *Botrytis cinerea* has been shown to increase tolerance to H_2_O_2_-derived oxidative stress and host pathogenicity while negatively affecting growth and development (Zhang et al. 2018).

Glutathione pathway components were among the most significantly altered in response to oxidative stress in both isolates, though to a greater extent in NRRL3357 which can be seen in the higher accumulation of oxidized glutathione, 5-oxoproline, and ophthalmate in NRRL3357 under stress which were not seen in AF13 (Figure 3). This pathway in conjunction with enzymes such as catalases and thioredoxin reductases and peroxidases serve as the primary means of redox homeostasis and oxidative stress alleviation for eukaryotes (Breitenbach et al. 2015). Glutathione metabolism has been previously linked to both development and aflatoxin production in *Aspergillus spp*. Huang et al. (2009) showed that treatment of *A. flavus* with an ethylene-producing compound resulted in increases in GSH/GSSH ratios, oxidative stress remediation, and significant reductions in aflatoxin biosynthetic gene expression and aflatoxin production. Reduced glutathione accumulation has also been associated with asexual and sexual development in *A. nidulans* thioredoxin A (*AnTrxA*) mutants with applied GSH resulting in restored conidiation and early induction of cleistothecia formation following long-term, low concentration application (Thon et al. 2007). Given this relationship between glutathione, development, and mycotoxin production, this mechanism may be lending to distinctive growth patterns and aflatoxin production levels in these isolates which represent diverse VCGs and mating types and warrants further investigation (Horn, 2007; Mehl and Cotty, 2010, 2013).

Sulfur-containing amino acids such as cysteine and methionine, and their derivatives were also differentially accumulated in response to oxidative stress (Figure 4). These compounds have antioxidant benefits, and also function in important signaling capacities. Taurine, an antioxidant compound (Jong et al. 2012), was shown to accumulate in both isolates under stress along with its immediate precursor 3-sulfo-L-alanine which may supplement other antioxidant pathways (Figure 4). The detected signaling compounds, SAM and MTA, are closely tied to polyamine biosynthesis which was also significantly regulated by oxidative stress. Polyamines such as putresine and spermidine differentially accumulated in this experiment (Figure 4), and have been found to function in oxidative stress responses either by scavenging ROS, inhibiting ROS-generating enzymes, or functioning in signal transduction to promote antioxidant mechanisms (Valdes-Santiago and Ruiz-Herrera, 2014). S-adenosylmethionine is required for the production of polyamines and MTA is produced from decarboxylated SAM by spermidine synthase and spermine synthase with accumulating MTA being able to inhibit these enzymes to prevent the generation of H_2_O_2_-derived oxidative stress due to polyamine back-conversion (Avila et al. 2004). Therefore, polyamine metabolism along with glutathione metabolism form a coordinated basis for regulating cellular redox potential in *A. flavus* in response to oxidative stress and may assist in coordination of both reproductive development and mycotoxin production.

Fatty acids were also significantly altered in accumulation in response to oxidative stress. This is particularly true for mono- and poly-unsaturated fatty acids which tended to be increased in abundance in NRRL3357 in response to stress, but not in AF13 (Figure 5). Unsaturated fatty acids have been found to be suitable substrates for aflatoxin production by *A. flavus* and *A. parasiticus*, and their byproducts have been shown to regulate aflatoxin production and development (Fanelli and Fabbri, 1989; Tsitsigiannis and Keller, 2007). For example, linoleic acid derivatives known as Psi factors have been shown to regulate both asexual and sexual sporulation in *A. flavus* and *A. nidulans* (Calvo et al. 1999), and oxylipins function in signaling for development, mycotoxin production, and host interactions (Affeldt et al. 2012; Fischer and Keller, 2016; Gao et al. 2009). In addition to signaling, free fatty acids also serve as important sources of energy, and can be catabolized to produce other macromolecules. Here, a majority of unsaturated lipids were depleted over time in control and stressed conditions in both isolates likely to provide energy and components for repairing and responding to oxidative stress (Supplemental File 1). These fatty acids are also important for maintaining membrane integrity and fluidity under environmental stress conditions. For example, dienoic fatty acids have been shown to function in preserving membrane fluidity in yeast under freezing and salt stresses (Rodriguez-Vargas et al. 2007).

Along with these major classes of metabolites, several cofactors and secondary metabolites were also differentially accumulated in response to stress (Supplemental File 1). Of particular interest were mevalonate and related terpenoid compounds which were increased in both isolates in response to oxidative stress. These compounds are precursors to some isoprenoid mycotoxins such as aflatrem whose biosynthetic genes have been found to be upregulated in response to oxidative stress in these isolates (Fountain et al. 2016a, 2016b). In addition, mevalonate and its derivatives have been shown to link the biosynthetic pathways for ergosterol and ornithine-derived siderophores, and interruption of this link results in reduced tolerance to oxidative stress, siderophore production, and virulence in *A. fumigatus* (Yasmin et al. 2011).

These compounds differentially accumulating in these isolates of *A. flavus* mirror those observed in other *Aspergillus spp.* such as *A. oryzae* (Singh et al. 2018) and provide potential insights into putative approaches to enhance host resistance under drought stress. We hypothesized that excessive ROS generated in drought sensitive host plants during drought stress may contribute to enhancing susceptibility to aflatoxin contamination (Fountain et al. 2014; 2015). In addition, the metabolic pathways employed by *A. flavus* in remediating oxidative stress seen in this study parallel those employed by host plants such as maize in countering drought stress in developing kernel tissues (Yang et al. 2018). Given this relationship, the manipulation of host tissue composition may be a viable approach to improve aflatoxin contamination resistance through two possible methodologies. The first method is biomarker selection employed in breeding programs (Fernandez et al. 2016). For aflatoxin mitigation, enhanced accumulation of antioxidant compounds in host plant tissues corresponding to those observed in *A. flavus* such as glutathione pathway components, polyamines, or simple sugar content could be selected for in conventional and molecular breeding programs. The second method is genetic engineering including both genome editing and transgenic approaches to manipulate the expression of host plant enzymes to modify kernel composition to reduce stress on infecting A. flavus under drought. These technologies could also be used to enhance host plant antioxidant potential through increased antioxidant enzyme expression, antioxidant compound production, or aflatoxin inhibitor production. This would also have the added potential benefit of reduced drought-related kernel abortion and filling reduction due to oxidative damage.

## Materials and Methods

### Isolate collection

The isolates used in this study were obtained as follows. AF13 was requested from Dr. Kenneth Damann, Department of Plant Pathology and Crop Physiology, Louisiana State University, Baton Rouge, LA. NRRL3357 was requested from the USDA National Culture Repository, Peoria, IL. All isolates were shipped on PDA and transferred to V8 agar as previously described (Fountain et al. 2018). Agar plugs containing fresh conidia were taken along the growing edge of the colonies and stored in sterile water and 20% (v/v) glycerol at 4 and −20°C, respectively, until used.

### Culture conditions and tissue collection

Isolate conidia suspensions were used to inoculate V8 agar plates, and were incubated at 37°C for 5 days. Conidia were then harvested using sterile 0.1% (v/v) Tween 20 and a sterile loop to make a fresh conidia suspension (~2.0 x 10^7^ conidia/mL) for use as inoculum. For each isolate, 100 µL of conidial suspension was then used to inoculate stationary liquid cultures of 50 mL yeast extract-sucrose medium (YES; 2% yeast extract, 1% sucrose) in 125mL Erlenmeyer flasks amended with H_2_O_2_ (3% stabilized solution) to a final concentration of either 0 or 15mM and a final conidia concentration of ~8.0 x 10^4^ conidia/mL. The flasks were plugged with sterile cotton and incubated at 30°C in the dark. Mycelial mats were then harvested for each isolate and H_2_O_2_ treatment at 4 and 7 days after inoculation (DAI). Five repeat cultures representing five biological replicates were harvested for each isolate, treatment, and time point. A detailed description of the experiment design can be found in Figure S1. Harvested mycelia mats were immediately flash frozen in liquid nitrogen and ground into a fine powder using sterile, chilled mortar and pestles. The ground tissue (~1g) was then transferred to a sterile 2.0mL microcentrifuge tube and stored at −80°C until use in metabolomics analysis.

### Metabolomic profiling

Collected and ground mycelia tissues were used for global, unbiased metabolomics by Metabolon (Morrisville, NC, USA) as described by Yang et al. (2018) and Lin et al. (2017). Briefly, 50 mg of tissue from each sample were prepared using an automated MicroLab STAR system (Hamilton, Reno, NV, USA) during which QC standards were added for downstream normalization. Metabolites and proteins were extracted in methanol in a GenoGrinder 2000 (Glen Mills, Clifton, NJ, USA) followed by centrifugation for metabolite isolation and protein separation. Each extract was then divided into 5 fractions and used for reverse phase (RP)/ultra-performance liquid chromatography (UPLC)-tandem mass spectrometry (MS/MS) with positive ion mode electrospray ionization (ESI), RP/UPLC-MS/MS with negative ion mode ESI, and HILIC/UPLC-MS/MS with negative ion mode ESI. One fraction from each extract was reserved as a backup. All methods employed either an ACQUITY UPLC (Waters, Milford, MA, USA) or a Q-Exactive High Resolution/Accuracy Mass Spectrometer with a heated electrospray ionization (HESI-II) source and an Orbitrap Mass Analyzer (ThermoFisher, Waltham, MA, USA). A detailed description of methods and procedure for data acquisition, metabolite acquisition, quantitation, and data analysis can be found in Supplemental File 2.

### Growth curve assay

A growth curve assay was performed for the isolates used for metabolomics analysis under H_2_O_2_-derived stress using a microtiter plate method as described by Meletiadis et al. (2001). Both isolates were cultured on V8 agar for 7 days at 30°C in the dark. Agar plugs were collected along the growing edge of the colonies and placed into amber bottles containing ~5.0 mL 0.1% (v/v) Tween 20 and gently shaken to suspend conidia. The concentration of each conidial suspension was measured using a hemocytometer, and used to prepare inoculum for each isolate with at two concentrations of 2.0 x 10^4^ conidia/mL as described by Meletiadis et al. (2001) and 8.0 x 10^4^ conidia/mL as used for the present metabolomics assay. A 96-well flat bottom microtiter plate was then prepared by filling each well with 100 µL of double strength YES medium (4% yeast extract, 2% sucrose) amended with 0, 20, 30, 40, 50, or 60 mM H_2_O_2_. For each inoculum, 100 µL was added to each the prepared wells resulting in a standard YES concentration and a final concentration of 0, 10, 15, 20, 25, or 30 mM H_2_O_2_. Three replicate wells were inoculated for each isolate and treatment combination. For non-inoculated wells, 100 µL of 0.1% Tween 20 was added in place of inoculum. The plate was sealed with optically-clear tape and incubated at 30°C in the dark without shaking in a Synergy HT plate reader (Biotek, Winooski, VT, USA). Optical density at 405 nm (OD_405_) was recorded every 15 min for 100 hr. The average of the OD_405_ for the non-inoculated wells was then subtracted from each measurement to remove background absorbance.

### Data analysis

Raw data obtained from UPLC-MS/MS analyses were peak-identified and QC corrected based on the Metabolon Laboratory Information Management System (LIMS) which contains identifying information for >4500 standard compounds. Quantitation and differential accumulation analyses were performed as described by Lawton et al. (2008), Lin et al. (2017), and Rao et al. (2014) using ArrayStudio and R (v3.4.0). Heatmaps and principal components analyses were performed using MultiExperiment Viewer (MeV, v4.9.0). Functional enrichment analyses were performed with Blast2GO (Conesa et al. 2005), and metabolic pathways were identified based on the KEGG database (Kanehisa and Goto, 2000). Pearson correlation analyses of the detected metabolites was performed using R (v3.4.0) and RStudio (v1.1.423). For the growth curve analysis, Gen3 software (Biotek) was then used to calculate the highest OD (OD_max_), and average time of initial detection at a defined threshold of OD_405_ = 0.2 (T_i_).

## Acknowledgements

We would like to thank Billy Wilson and Hui Wang for technical assistance in the laboratory. This work is partially supported by the U.S. Department of Agriculture Agricultural Research Service (USDA-ARS), USDA National Institute for Food and Agriculture (USDA-NIFA), the Georgia Agricultural Commodity Commission for Corn, the National Corn Growers Association Aflatoxin Mitigation Center of Excellence (AMCOE), the Georgia Peanut Commission, and The Peanut Foundation. Mention of trade names or commercial products in this publication is solely for the purpose of providing specific information and does not imply recommendation or endorsement by the USDA. The USDA is an equal opportunity employer and provider.

## Author Contributions

JCF performed the culture experiments and data analyses, and wrote the manuscript. LY, MKP, and PB assisted in data analysis and in project discussions. DA performed the metabolomics experiment and assisted in data analysis. SC, RCK, and RKV contributed to project discussions and assisted with revision of the manuscript. BG conceived the project, planned, secured extramural funds, and revised and submitted manuscript.

## Supplemental Figure Legends

**Figure S1**. Metabolomics experiment design. Two isolates of *Aspergillus flavus*, AF13 (highly aflatoxigenic and oxidative stress tolerant) and NRRL3357 (moderate to highly aflatoxigenic and moderately oxidative stress tolerant), were grown in yeast extract sucrose (YES) medium supplemented with either 0 or 15 mM H_2_O_2_. Samples were collected at 4 and 7 days after inoculation (DAI). Five biological replicates (n = 5, N = 40) were performed for each isolate, treatment, and time point combination. Statistical comparisons are indicated by the colored arrows with blue indicating oxidative stress effect comparisons, red indicating time effects, and green indicating isolate/genotype effects.

